# Mutational signatures and heterogeneous host response revealed via large-scale characterization of SARS-CoV-2 genomic diversity

**DOI:** 10.1101/2020.07.06.189944

**Authors:** Alex Graudenzi, Davide Maspero, Fabrizio Angaroni, Rocco Piazza, Daniele Ramazzotti

## Abstract

To dissect the mechanisms underlying the inflation of variants in the SARS-CoV-2 genome, we present one of the largest up-to-date analyses of intra-host genomic diversity, which reveals that most samples present heterogeneous genomic architectures, due to the interplay between host-related mutational processes and transmission dynamics.

The deconvolution of the set of intra-host minor variants unveils the existence of non overlapping mutational signatures related to specific nucleotide substitutions, which prove that distinct hosts respond differently to SARS-CoV-2 infections, and which are likely ruled by APOBEC, Reactive Oxygen Species (ROS) and ADAR.

Thanks to a corrected-for-signatures *dN/dS* analysis we demonstrate that the mutational processes underlying such signatures are affected by purifying selection, with important exceptions. In fact, several mutations linked to low-rate mutational processes appear to transit to clonality in the population, eventually leading to the definition of new clonal genotypes and to a statistically significant increase of overall genomic diversity.

Importantly, the analysis of the phylogenetic model shows the presence of multiple homoplasies, due to mutational hotspots, phantom mutations or positive selection, and supports the hypothesis of transmission of minor variants during infections. Overall, the results of this study pave the way for the integrated characterization of intra-host genomic diversity and clinical outcome of SARS-CoV-2 hosts.

## Introduction

The COVID-19 pandemic has currently affected 216 countries and territories worldwide with ≈ 70 million people being infected, while the number of casualties has reached the impressive number of ≈ 1.6 million (World Health Organization (WHO) (2020), update 15 December 2020). The origin and the main features of SARS-CoV-2 evolution have been investigated (Zhou et al., 2020b; Wu et al., 2020; Andersen et al., 2020; Xiao et al., 2020; Deng et al., 2020), also due to the impressive amount of consensus viral sequences included in public databases, such as GISAID (Shu and McCauley, 2017). However, only a few currently available datasets include raw sequencing data, which are necessary to quantify intra-host genomic variability.

Due to the combination of high error and replication rates of viral polymerase, subpopulations of viruses with distinct genotypes, also known as viral quasispecies (Domingo et al., 1985), usually coexist within single hosts. Such heterogeneous mixtures are supposed to underlie most of the adaptive potential of RNA viruses to internal and external selection phenomena, which are related, e.g., to the interaction with the host’s immune system or to the response to antiviral agents. For instance, it was hypothesized that intra-host heterogeneity may be correlated with prognosis and clinical outcome (Novella et al., 1995; Domingo et al., 2012). Furthermore, even if the modes of transmission of intra-host variants in the population are still elusive, one may hypothesize that, in certain circumstances, infections allow such variants to spread, sometimes inducing significant changes in their frequency (Lythgoe et al., 2020).

In particular, several studies on SARS-CoV-2 support the presence of intra-host genomic diversity in clinical samples and primary isolates (Ramazzotti et al., 2020; Shen et al., 2020; Wölfel et al., 2020; Capobianchi et al., 2020; Rose et al., 2020; Lu et al., 2020; Lythgoe et al., 2020; Seemann et al., 2020; Popa et al., 2020), whereas similar results were obtained on SARS-CoV (Xu et al., 2004), MERS (Park et al., 2016), EBOLA (Ni et al., 2016) and H1N1 influenza (Poon et al., 2016). We here present one of the the largest up-to-date studies on intra-host genomic diversity of SARS-CoV-2, based on a large dataset including 1133 high-quality samples for which raw sequencing data are available (NCBI BioProject: `PRJNA645906`). The results were validated on 4 independent datasets including a total of 953 samples (`PRJNA625551, PRJNA633948, PRJNA636748, PRJNA647529`; see the Validation section).

Our analysis shows that ≈ 15% of the SARS-CoV-2 genome has already mutated in at least one sample, including ≈ 1% of positions exhibiting multiple mutations. The large majority of samples shows a heterogeneous intra-host genomic composition, with 892 out of 1133 samples (≈ 79%) exhibiting at least one low frequency variant (variant frequency, VF > 5% and ≤ 90%, named *minor variants* or *iSNVs*), 171 samples more than 5 and 101 samples more than 10. Importantly, several variants are observed as clonal (VF > 90%) in certain samples and at a low frequency in others, demonstrating that transition to clonality might be due not only to functional selection shifts, but also to complex transmission dynamics involving bottlenecks and founder effects (Domingo et al., 2012).

Strikingly, our analysis allowed to identify three non-overlapping *mutational signatures*, i.e., specific distributions of nucleotide substitutions,n which are observed in distinct mixtures and with significantly different intensity in three well-separated *clusters* of samples, suggesting the presence of host-related mutational processes. One might hypothesize that such processes are related to the interaction of the virus with the host’s immune system, and might pave the way for a better understanding of the molecular mechanisms underlying different clinical outcomes.

In particular, the first signature is dominated by C>T:G>A substitution and is likely related to APOBEC activity, the second signature is mostly characterized by G>T:C>A substitution and might be associated to ROS-related processes, while a third signature is predominantly associated to A>G:T>C substitution, which is usually imputed to ADAR activity. A corrected-for-signatures version of the *dN/dS* analysis would suggest that, as expected, the three signatures are affected by mild purifying selection in the population, yet with some exceptions that would suggest the existence of positively selected genomic regions. Furthermore, a certain proportion of samples of two signature-based clusters, mostly associated to APOBEC and ROS appear to be hypermutated (up to 87 minor variants detected in a single host), whereas this effect is mitigated for the remaining cluster, dominated by ADAR-related processes.

Finally, the analysis of the phylogenetic model, obtained from the profiles of clonal variants via VERSO (Ramazzotti et al., 2020), allowed us to assess how many minor variants are either detected in single samples, in multiple samples of the same clade, or in multiple samples of independent clades (i.e., *homoplasies*). Strikingly, an approximately monotonic decrease of the median variant frequency is observed with respect to the number of clades in which minor variants are observed: minor variants detected in single clades exhibit the largest (median) variant frequency, as opposed to variants shared in multiple clades, which display a progressively lower variant frequency.

On the one hand, this result supports the hypothesis of transmission of minor variants during infections, and of the concurrent existence of bottleneck effects (Gutierrez et al., 2012; Domingo et al., 2012). On the other hand, the significant number of minor variants observed at a low frequency in multiple clades would suggest the presence of mutational hotspots and of phantom mutations related to sequencing artifacts (Bandelt et al., 2002).

## Results

### Mutational landscape of SARS-CoV-2 from variant frequency profiles of 1133 samples – Dataset #1

We performed variant calling from Amplicon raw sequencing data of 1188 samples from the NCBI BioProject `PRJNA645906` and by aligning sequences to reference genome SARS-CoV-2-ANC, which is a likely ancestral SARS-CoV-2 genome (Ramazzotti et al., 2020). The mutational profiles of 1133 high-quality samples selected after quality check was analyzed in-depth (see Methods).

In detail, 4677 distinct single-nucleotide variants (SNVs, identified by genome location and nucleotide substitution) were detected in the dataset, for a total of 19663 non-zero entries of the *variant frequency* (VF) matrix (see Methods for further details; the VF profiles of all samples are included in Supplementary File S1; see Fig. 1H for a graphical representation of an example dataset). In particular, in our analysis we consider any SNV detected in any given sample as *clonal*, if its VF is > 90% and as *minor* if its VF is > 5% and ≤ 90%.

**Figure 1:**
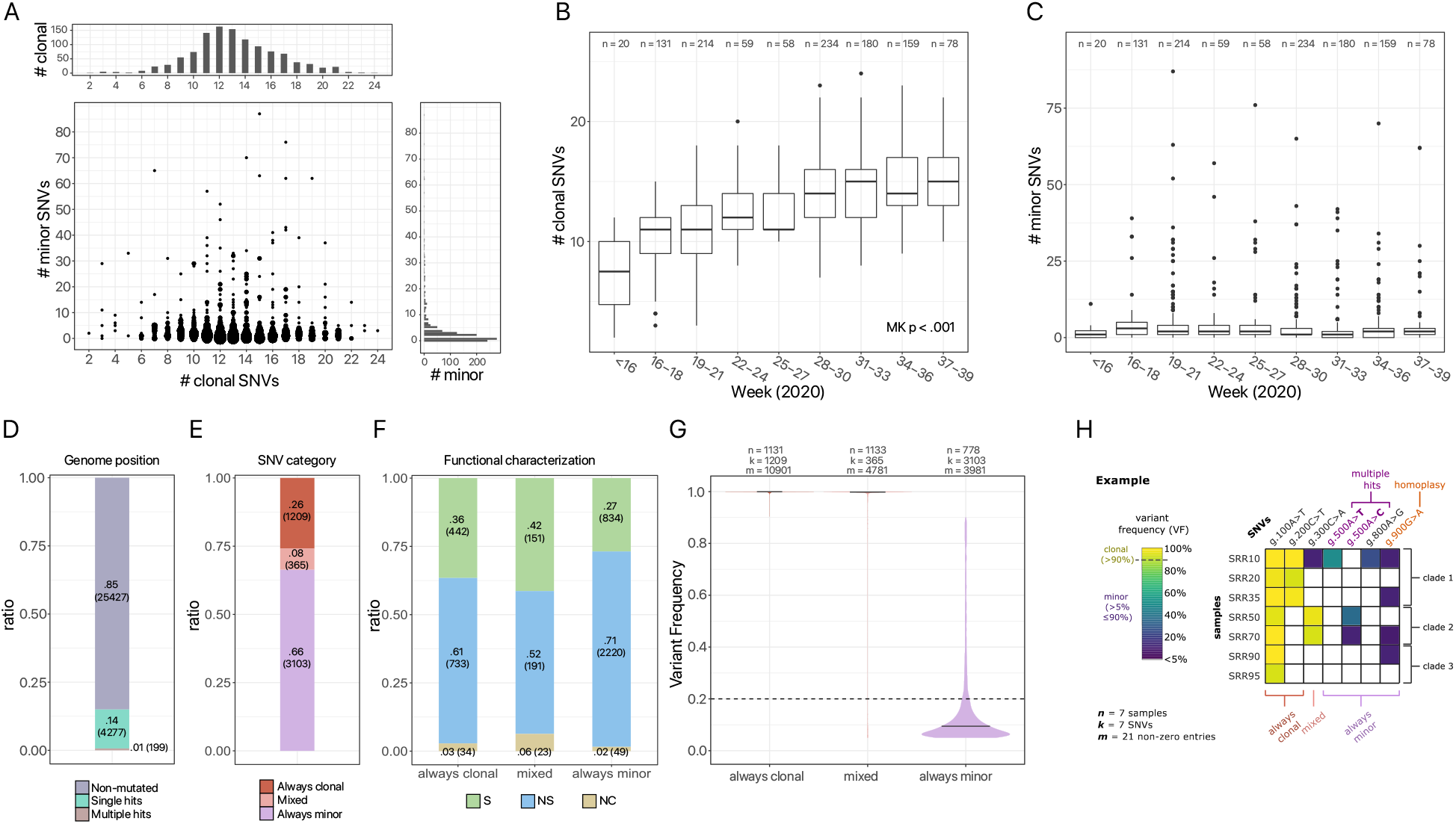
Mutational landscape of 1133 SARS-CoV-2 samples – Dataset #1 (PRJNA645906) (**A**) Scatter-plot displaying the number of clonal (VF > 90%) and minor (VF > 5% and ≤ 90%) variants for 1133 samples of Dataset #1 (node size proportional to the number of samples). (**B**) Box-plots returning the distribution of the number of clonal and (**C**) minor variants, obtained by grouping samples according to collection date (weeks, 2020; Mann-Kendall trend test p-value also shown). *n* returns the number of samples in each group. (**D**) Bar-plot returning the proportion of sites of the SARS-CoV-2 genome that are either non-mutated, mutated with a unique SNV, or mutated with multiple SNVs. (**E**) Stacked bar-plots returning the proportion of SNVs detected as always clonal, mixed or always minor. (**F**) The ratio of synonymous (S), non-synonymous (NS) and non-coding (NC) mutations, for each category. (**G**) Violin plots returning the distribution of VF of all SNVs (*n* returns the number of samples, *k* the number of distinct SNVs, *m* the number of non-zero entries of the VF matrix). (**H**) Graphical representation of an example dataset.

The distribution of the number of minor and clonal variants observed in each sample (Fig. 1A) unveils an approximately normal distribution of clonal variants (median = 13, mean = 13.2 and max = 24). Minor variants are detected in ≈ 78.7% of the samples and show a long-tail distribution (median = 2, mean = 4.16 and max = 87). 109 samples (≈ 9.6% of the dataset) display a number of minor variants ≥ 10, up to a maximum of 87.

Interestingly, we observe a statistically significant increase of genomic diversity on clonal variants with respect to collection week (Mann-Kendall test for trend on median number of clonal variants *p* < 0.001, Fig. 1B), due to the accumulation of clonal variants in the population, and which confirms recent findings (Li et al., 2020; Shen et al., 2020; Ramazzotti et al., 2020), whereas, as expected, this phenomenon is less evident for minor variants (Fig. 1C). This aspect is further investigated in the following and hints at the interplay involving the evolutionary dynamics within hosts and the transmission among hosts, which differently affects clonal and minor variants (Chan et al., 2013).

#### Evidence of transition to clonality

We further categorize each detected SNV as: (*i*) *always clonal*, if clonal in all samples in which it is detected, (*ii*) *always minor*, if minor in all samples in which it is detected, (*iii*) *mixed*, if observed as clonal in at least one sample and as minor in at least another sample.

4677 single-nucleotide variants were detected on 4476 distinct genome sites (≈ 14.9% of the SARS-CoV-2 genome), of which 199 sites (≈ 0.6% of the genome) display multiple nucleotide substitutions (see Fig. 1D). This suggests that the proportion of mutated genomic sites might be considerably higher in the overall population, especially if considering minor variants. Overall, 25.8%, 7.8% and 66.3% SNVs are detected as always clonal, mixed and always minor, and are mostly non-synonymous (see Fig. 1E–F).

The analysis of the VF distribution (Fig. 1G) unveils an impressive scarcity of variants showing VF in the middle-range, i.e., between 20% and 90%, for all categories. This phenomenon is likely due to transmission *bottlenecks*, which tend to purify low-frequency variants in the population. Nonetheless, both mixed and always minor variants display broad VF spectra, an aspect that is particularly relevant for the former category. In this respect, 24.4% of all mixed variants (89 on 365) never display a VF ≤ 20%: one may hypothesize that such variants are indeed *transiting to clonality* in the population, because either positively selected, as a result of the strong immunologic pressure within human hosts (Lucas et al., 2001), or because affected by transmission phenomena involving *founder effects, bottlenecks* and *stochastic fluctuations* (Gutierrez et al., 2012; Domingo et al., 2012). Conversely, one might hypothesize that most remaining mixed variants may result from random mutations hitting positions of SNVs that are already present as clonal in the population.

Furthermore, the distribution of SNVs with respect to each region of the genome in Fig. 2A demonstrate that mutations are approximately uniformly distributed across the genome (see also Supplementary Fig.s S2-S4). Overall, this analysis provides one of the first large-scale quantifications of transition to clonality in SARS-CoV-2 and might serve to intercept variants possibly involved in functional modifications, bottlenecks or founder effects.

**Figure 2:**
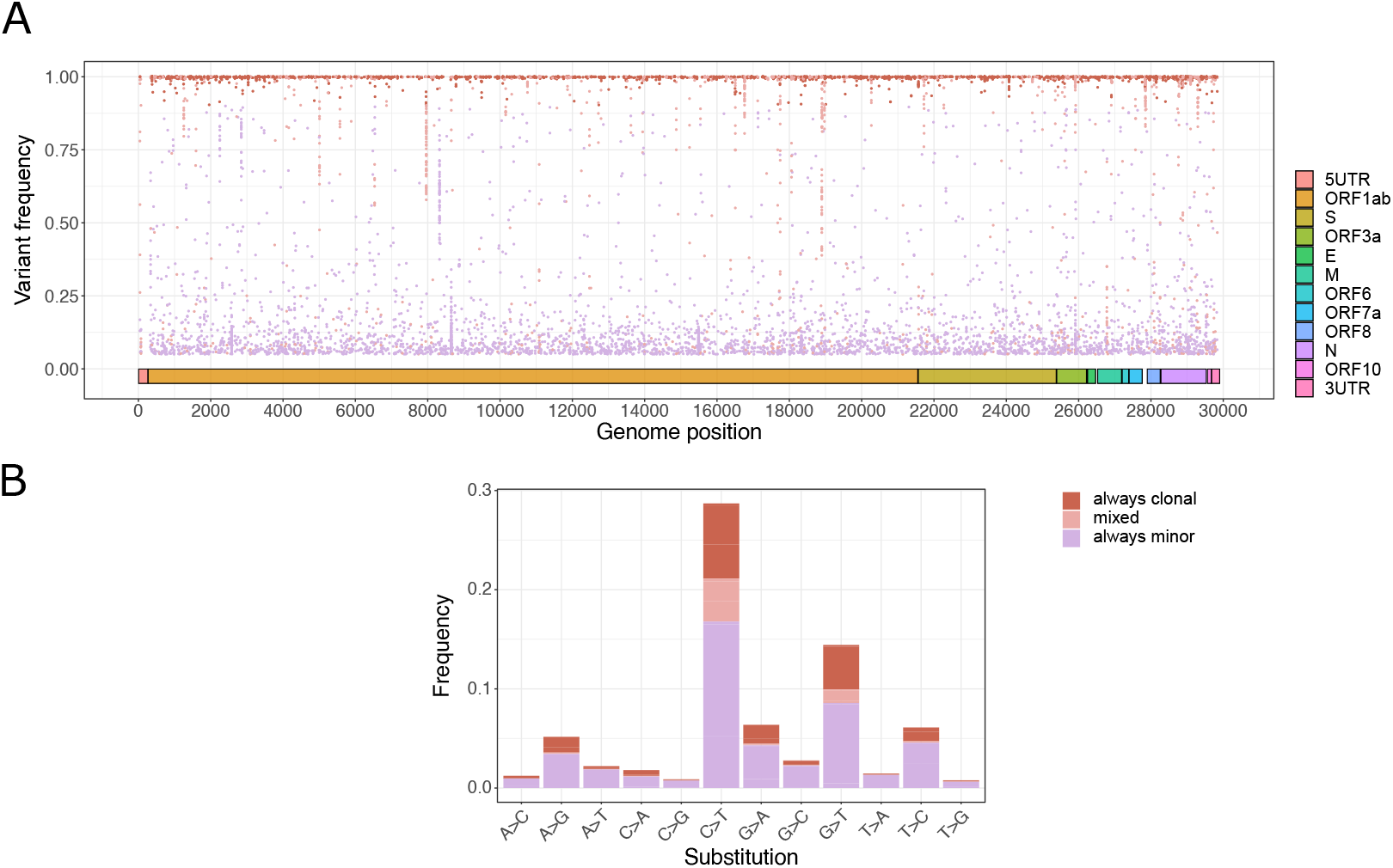
Characterization of SNVs detected on the SARS-CoV-2 genome. (**A**) Scatter-plot returning the genome location and the VF of all SNVs detected in the dataset, colored according to category. (**B**) Stacked bar-plots returning the normalized substitution proportion of all SNVs detected in at least one sample of the dataset, with respect to all 12 possible nucleotide substitutions, grouped by variant type.

#### De novo decomposition of SARS-CoV-2 mutational signatures

In order to investigate the existence of mutational processes related to the interaction between the host and the SARS-CoV-2 virus, we analyzed the distribution of nucleotide substitutions for all SNVs detected in the dataset. In Fig. 2B one can see the proportion of SNVs for each of the 12 nucleotide substitution types (e.g., number of C>T’s) over the total number of nucleotides present in the reference genome for each substitution type (e.g., number of C’s).

Certain substitutions present a significantly higher normalized abundance, confirming recent findings on distinct cohorts (Simmonds, 2020; Di Giorgio et al., 2020; Popa et al., 2020). In particular, C>T substitutions are observed in ≈ 28% of all C nucleotides in the SARS-CoV-2 genome; G>T’s in ≈ 15% of all G’s, T>C’s in ≈ 7% of all T’s, and A>G’s in ≈ 5% of all A’s. Although, traditionally, a 12-substitution pattern has been used in order to report mutations occurring in single-stranded genomes, we reasoned that, owing to the intrinsically double-stranded nature of the viral life-cycle (i.e., a mutation occurring on a *plus* strand can be transferred on the *minus* strand by RdRP and vice versa) it is sound to consider a total of 6 substitution classes (obtained by merging equivalent substitutions in complementary strands) to investigate the possible presence of viral mutational signatures (Alexandrov et al., 2013). Clonal variants were not considered in the next analyses, to focus on SNVs likely related to host-specific mutational processes and by excluding variants presumably transmitted during infection events.

In particular, in order to identify and characterize the mutational processes underlying the emergence of SARS-CoV-2 variants with a statistically grounded approach, we applied a Non-Negative Matrix Factorization (NMF) approach (Brunet et al., 2004) and standard metrics to determine the optimal rank (see Methods). In particular, we analyzed the mutational profiles of 150 samples exhibiting at least 6 always minor variants (on 1133 total samples), to ensure a sufficient sampling of the distributions.

Strikingly, 3 distinct and non-overlapping mutational signatures are found and explain 96.5% of the variance in the data (Fig. 3A and Supplementary Fig. S3; Cophenetic correlation coefficient = 0.998, Cosine similarity between predictions and observations = 0.973, harmonic mean p-value of the one-sided Mann-Whitney U test on bootstrap re-sampling < 0.01 for all signatures, see Methods). In particular, signature *S*#1 is predominantly related to substitution C>T:G>A (81.2%), signature *S*#2 to substitution C>A:G>T (77.7%), while signature *S*#3 is dominated by substitutions T>C:A>G (60.7%) and T>A:A>T (23.6%).

**Figure 3:**
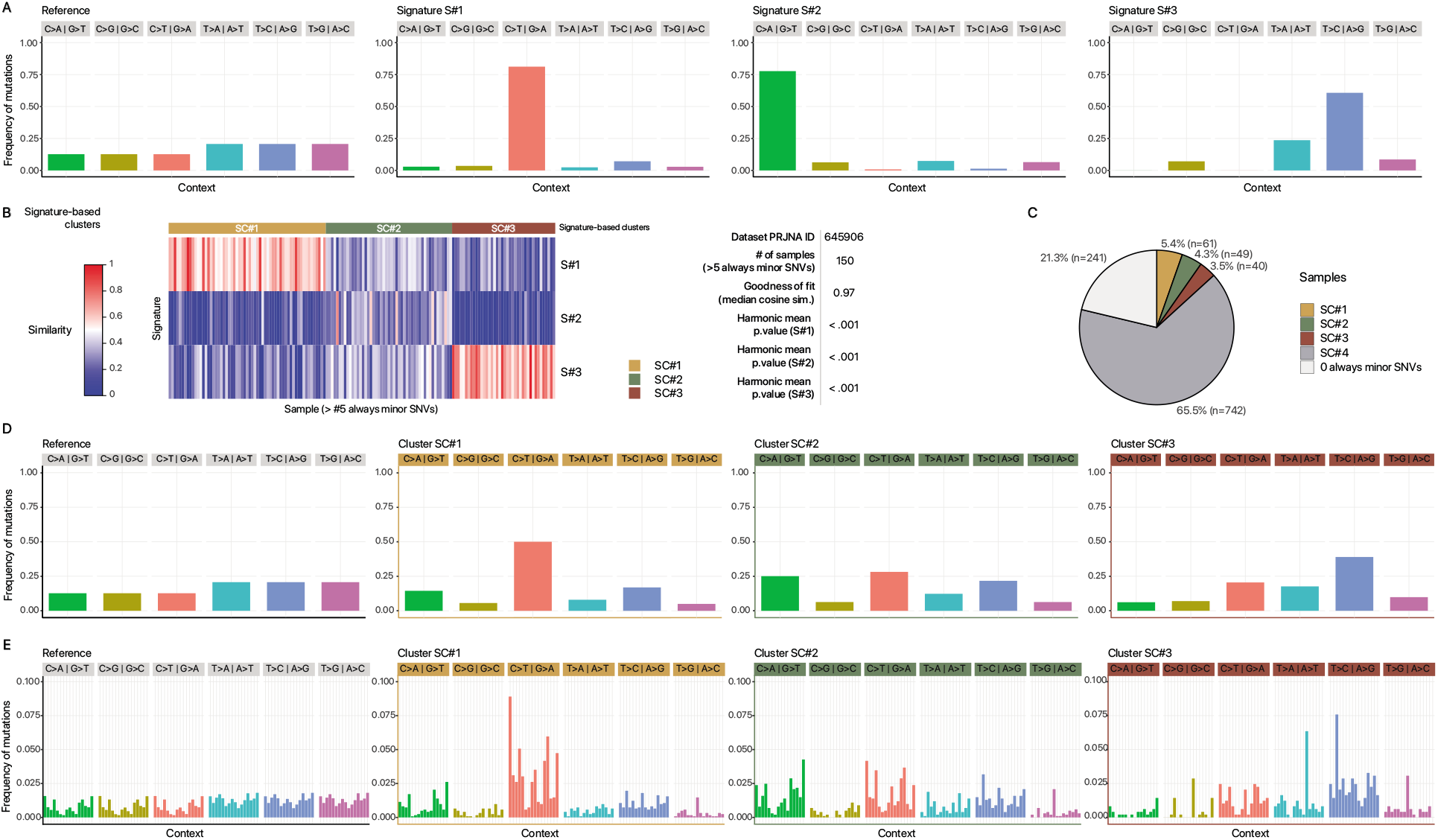
Mutational signatures of SARS-CoV-2. (**A**) The nucleotide class distribution in SARS-CoV-2-ANC reference genome (Ramazzotti et al., 2020) and for the 3 SARS-CoV-2 mutational signatures retrieved via NMF on 6 substitution classes are shown. (**B**) Heatmap returning the clustering of 150 samples with ? 6 always minor variants (≈ 13% of the dataset), computed via k-means on the low-rank latent NMF matrix. The goodness of fit in terms of median cosine similarity between observations and predictions, and the harmonic mean p-value of the one-sided Mann-Whitney U test on bootstrap re-sampling, are shown for all signatures, see Methods). (**C**) Pie-chart returning the proportion of samples in the three signature-based clusters, plus a fourth cluster SC#4 including all samples with ≥ 1 and < 6 always minor variants, and the group of samples with 0 always minor SNVs. (**D**) Categorical normalized cumulative VF distribution of all SNVs detected in each signature-based cluster, with respect to 6 substitution classes and to (**E**) 96 trinucleotide contexts, as compared to the theoretical distribution in SARS-CoV-2-ANC reference genome (left).

#### Characterization of mutational signatures of SARS-CoV-2

Signature S#1 is related to C>T:G>A substitution, which was often associated to APOBEC (*Apolipoprotein B mRNA Editing Catalytic Polypeptide-like*), i.e., a cytidine deaminase involved in the inhibition of several viruses and retrotransposons (Sharma et al., 2015). An insurgence of APOBEC-related mutations was observed in other coronaviruses shortly after spillover (Woo et al., 2007) and it was recently hypothesized that APOBEC-like editing processes might have a role in the response of the host to SARS-CoV-2 (Simmonds, 2020).

As specified above, a mutational process occurring on single-stranded RNA with a given pattern, e.g., C>T, could occur as a C>T mutation on the *plus* reference strand, but could similarly occur on the *minus* strand, again as a C>T substitution. However, C>T events originally occurring in the *minus* strand would be recorded as G>A owing to the mapping of the mutational event as a reverse-complement on the *plus* reference genome. Starting from these considerations and hypothesizing that the C>T:G>A substitution is mediated by APOBEC, which operates on singlestranded RNA and is similarly active on both strands, the analysis of the C>T / G>A ratio (or, more generally, of a *plus/minus* substitution ratio) should give an accurate measurement of the molar ratio between the two viral strands inside the infected cells.

In our case, by comparing the proportion of substitutions of all minor variants detected in the dataset, the ratio C>T / G>A is 5.1 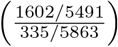. This result allows us to hypothesize that *plus* and *minus* viral strands of the SARS-CoV-2 genome are present in infected cells with a molar ratio in strong favor of the *plus* strand, and are consistent with the expected activity of APOBEC on single-stranded RNA. Further experimental analyses will be required to confirm this hypothesis.

The second signature S#2 is predominantly characterized by substitution C>A:G>T, whose origin is however still obscure. To gain insight into the mechanisms responsible for its onset, also in this case we analyzed the C>A and G>T substitution frequency, which revealed a strong disproportion in favor of the latter: the ratio G>T / C>A is 9.5 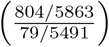. Overall, this results suggests that, in this case, the G>T substitution is the active mutational process.

In this respect, one might hypothesize a role for *Reactive Oxygen Species* (ROS) as mutagenic agent underlying this signature, as observed for instance in clonal cancer evolution (Alexandrov et al., 2020). ROS are extremely reactive species formed by the partial reduction of oxygen. A large number of ROS-mediated DNA modifications have already been identified; in particular however, guanine is extremely vulnerable to ROS, because of its low redox potential (David et al., 2007). ROS activity on guanine causes its oxidation to 7,8-dihydro-8-oxo-2’-deoxyguanine (oxoguanine). Notably: (*i*) oxoguanine can pair with adenine, ultimately causing G>T transversions, and (*ii*) ROS are able to operate on single-stranded RNA, therefore their mutational process closely resembles the C>A:G>T pattern we see in signature S#2. Thus, it is sound to hypothesize that the C>A:G>T substitution is generated by ROS, whose production is triggered upon infection, in line with several reports indicating that a strong ROS burst is often triggered during the early phases of several viral infections (Molteni et al., 2014; Reshi et al., 2014).

Finally, signature S#3 is primarily characterized by A>G:T>C substitution, which is typically imputed to the ADAR deaminase mutational process (Nishikura, 2010). ADAR targets adenosine nucleotides, causing deamination of the adenine to inosine, which is structurally similar to guanine, ultimately leading to an A>G substitution. Unlike APOBEC, ADAR targets double-stranded RNA, hence it is active only on *plus/minus* RNA dimers. In line with this mechanism and in sharp contrast with APOBEC, A>G’s and the equivalent T>C’s show a similar prevalence: the ratio A>G / T>C is 0.81 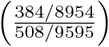. This supports the notion that the A>G:T>C mutational process is exquisitely selective for double-stranded RNA, where it can similarly targets adenines present on both strands.

#### Identification of signature-based clusters

We then clustered the 150 samples with at least 6 always minor mutations (on 1133 total samples), by applying k-means on the normalized low-rank latent NMF matrix and employing standard heuristics to determine the optimal number of clusters (see Methods). As a result, 3 signature-based clusters (SC#1, SC#2 and SC#3) are retrieved, including 61, 49 and 40 samples respectively (see Fig. 3B).

Remarkably, clusters SC#1 and SC#3 are characterized by distinctive signatures, S#1 (dominated by substitution C>T:G>A) and S#3 (T>C:A>G and T>A:A>T), respectively, whereas cluster SC#2 is characterized by a mixtures of all three signatures. In particular, the samples of the distinct clusters display dissimilar categorical VF distributions (see Fig. 3D), pointing at the existence of different host-related mutational processes. We here recall that samples with a number of always minor variants between 1 and 5 (742 samples, 65.5%) cannot be reliably associated to signature-based clusters, due to the low number of SNVs. For this reason, such samples were considered separately in the analysis and were labeled as cluster SC#4 from now on (Fig. 3C).

Importantly, by computing the categorical VF distribution of all minor SNVs with respect to all 96 trinucleotide contexts (i.e., by considering flanking bases), one can notice that clusters SC#1 and SC#2 display profiles that resemble that of the theoretical substitution distribution of the reference genome, thus suggesting that, in such cases, the host-related mutational processes are likely independent from flanking bases. Conversely, SC#3 displays a distribution of T substitutions with prevalent peaks in certain contexts and, especially, in G[T>A]G, A[T>C]G and C[T>G]T.

We finally note that, due to the possible transmission of minor variants among hosts during infections (see above), signature-based clusters might include both samples with host-related mutational processes and samples with minor variants herited from infecting hosts.

#### Characterization of signature-based clusters

We analyzed in-depth the intra-host genomic diversity of the samples of the 4 different signature-based clusters. As a first noteworthy result, while the distributions of the number of clonal variants are significantly alike across clusters (Kolmogorov-Smirnov, KS test *p* > 0.20 for 6/6 pairwise comparisons; see Fig. 4A and Supplementary File S2), clusters SC#1 and SC#2 display a similar distribution of minor variants (KS test *p* = 0.86), but significantly different distributions from the remaining clusters (KS *p* < 0.05 for all remaining pairwise comparisons; see Fig. 4B and Supplementary File S2). The relative proportion of substitution types for the samples of each signature-based cluster can be found in Fig. 4D.

**Figure 4:**
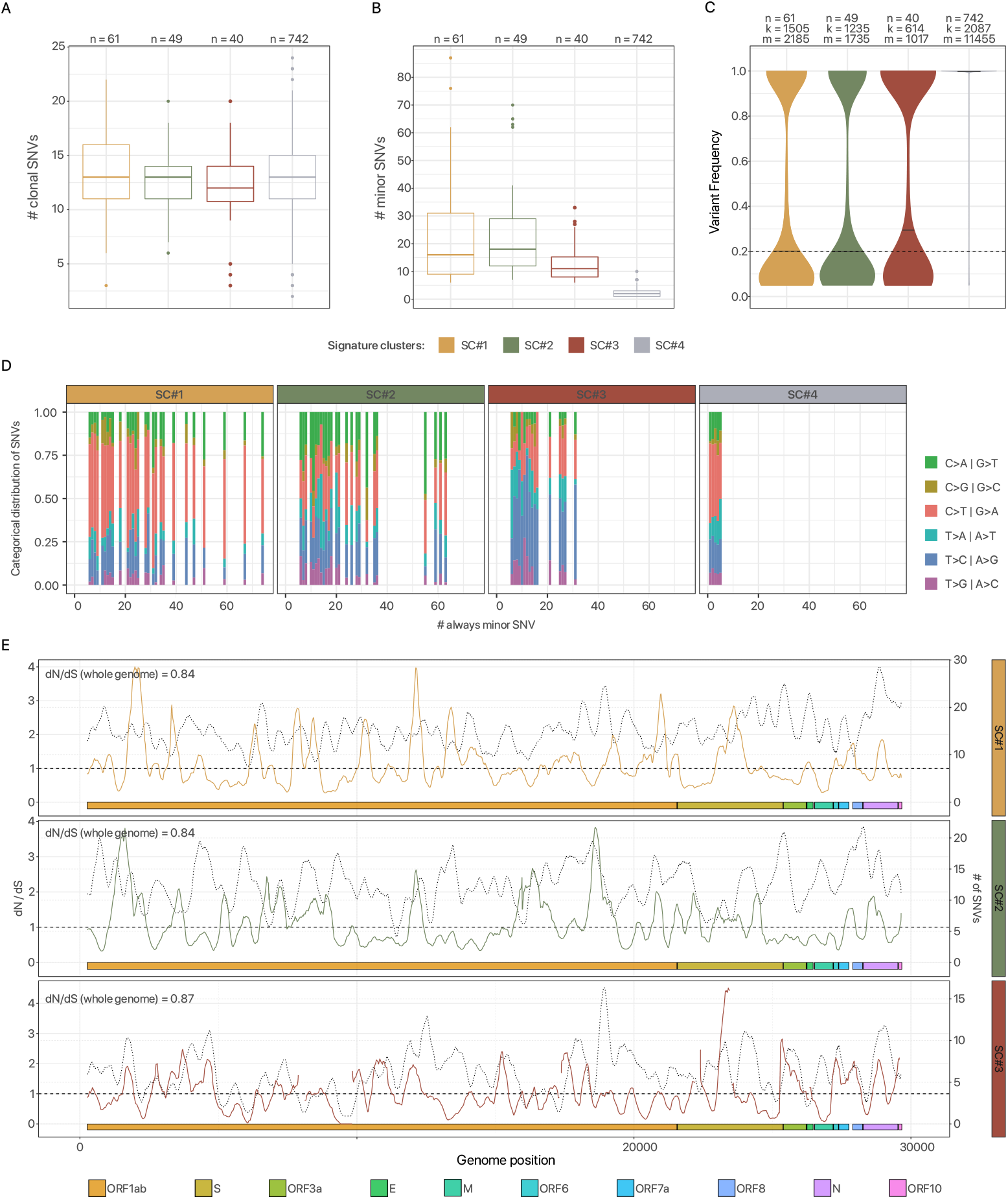
Characterization of signature-based clusters of SARS-CoV-2 samples. (**A**) Distribution of the number of clonal variants with respect to the 4 signature-based clusters described in the text. (**B**) Distribution of the number of minor variants for the 4 signature-based clusters. (**C**) Violin plots returning the VF distribution with respect to signature-based clusters (*n* returns the number of samples, *k* the number of distinct SNVs, *m* the number of non-zero entries of the VF matrix). (**D**) (Average) proportion of substitution classes of always minor variants for all the samples included in the 4 signature-based clusters, grouped and sorted by the number of minor SNVs (e.g., at position 10 of the x-axis one can find the average proportion of substitution classes for all samples with 10 minor SNVs). (**E**) Corrected-for-signatures *dN/dS* ratio plot, as computed by normalizing the ratio on cluster substitution distribution, on a 300-base sliding window, with respect to signature-based clusters (see Methods). The superimposed dotted line returns the mutational density in each window (rightmost y axis).

In particular, clusters SC#1 and SC#2 are characterized by a significantly higher number of minor variants (median 16 and 18, mean 22.3 and 22.6, max 87 and 70, for SC#1 and SC#2, respectively). Accordingly, both clusters include a certain proportion of highly mutated samples (with ≥ 10 minor variants), 44 on 61 and 41 on 49 for SC#1 and SC#2, respectively. This result supports the existence of highly active mutational processes and is consistent with the hypothesis of processes related to APOBEC and ROS. Conversely, cluster SC#3 displays a much lower number of minor variants (median 11, mean 13.2, max 33; 23 samples on 40 with ≥ 10 variants). This finding hints at the existence of milder spontaneous mutational processes related to ADAR.

Interestingly, the VF distribution for all SNVs highlights a remarkable similarity among signature clusters SC#1, SC#2 and SC#3, with the large majority of variants found either at a high or a low frequency, whereas, by construction, SC#4 is dominated by clonal variants (Fig. 4C). Moreover, only minor differences are observed in the distribution of substitutions with respect to SARS-CoV-2 Open Reading Frames (ORFs) (Supplementary Fig. S4).

Overall, these results reinforce the hypothesis of distinct mutational processes active in different hosts. When clinical data would be available in combination to sequencing data, this will allow to assess the correlation with clinical outcomes.

#### Evidence of purifying selection against signature-related mutagenic processes

To investigate the evolutionary dynamics of SARS-CoV-2, we implemented a corrected-for-signatures version of the *dN/dS* ratio analysis, i.e., obtained by normalizing the S/NS rate with respect to the theoretical distribution of substitutions detected in each cluster, as suggested in a different context in Van den Eynden and Larsson (2017).

Interestingly, the corrected *dN/dS* ratio computed on the genome coding regions (i.e., = 29133 basis) is equal to 0.84, 0.84 and 0.87, for the three signature-based clusters, respectively, and suggests the existence of purifying selection for all signature-related mutational processes.

We refined the analysis via a 300-base sliding window approach. On the one hand, the analysis of the mutational density confirms that the large majority of variants is indeed observed in purified regions of the genome. On the other hand, however, the variation of the corrected *dN/dS* ratio across the genome shows that some regions exhibit a ratio significantly larger than 1. This phenomenon, which is particularly evident in signature-based clusters SC#1 and SC#2, hints at possible positive selection processes affecting specific genomic regions and deserves further investigations.

#### Phylogenomic model of SARS-CoV-2 reveals transmission of minor variants and homoplasies

We employed VERSO (Ramazzotti et al., 2020) to reconstruct a robust phylogeny of samples from the binarized VF profiles of the 28 clonal variants (VF > 90%) detected in at least 3% of the dataset. In Fig. 5A one can see the output phylogenetic tree, which describes the existence of 23 clades and in which samples with identical corrected clonal genotype are grouped in polytomies (see Fig. 5B and the Methods section for further details). The mapping between clonal genotype labels and the lineage dynamic nomenclature proposed in Rambaut et al. (2020) and generated via pangolin 2.0 (O’Toole et al., 2020) is included in Supplementary File S3.

**Figure 5:**
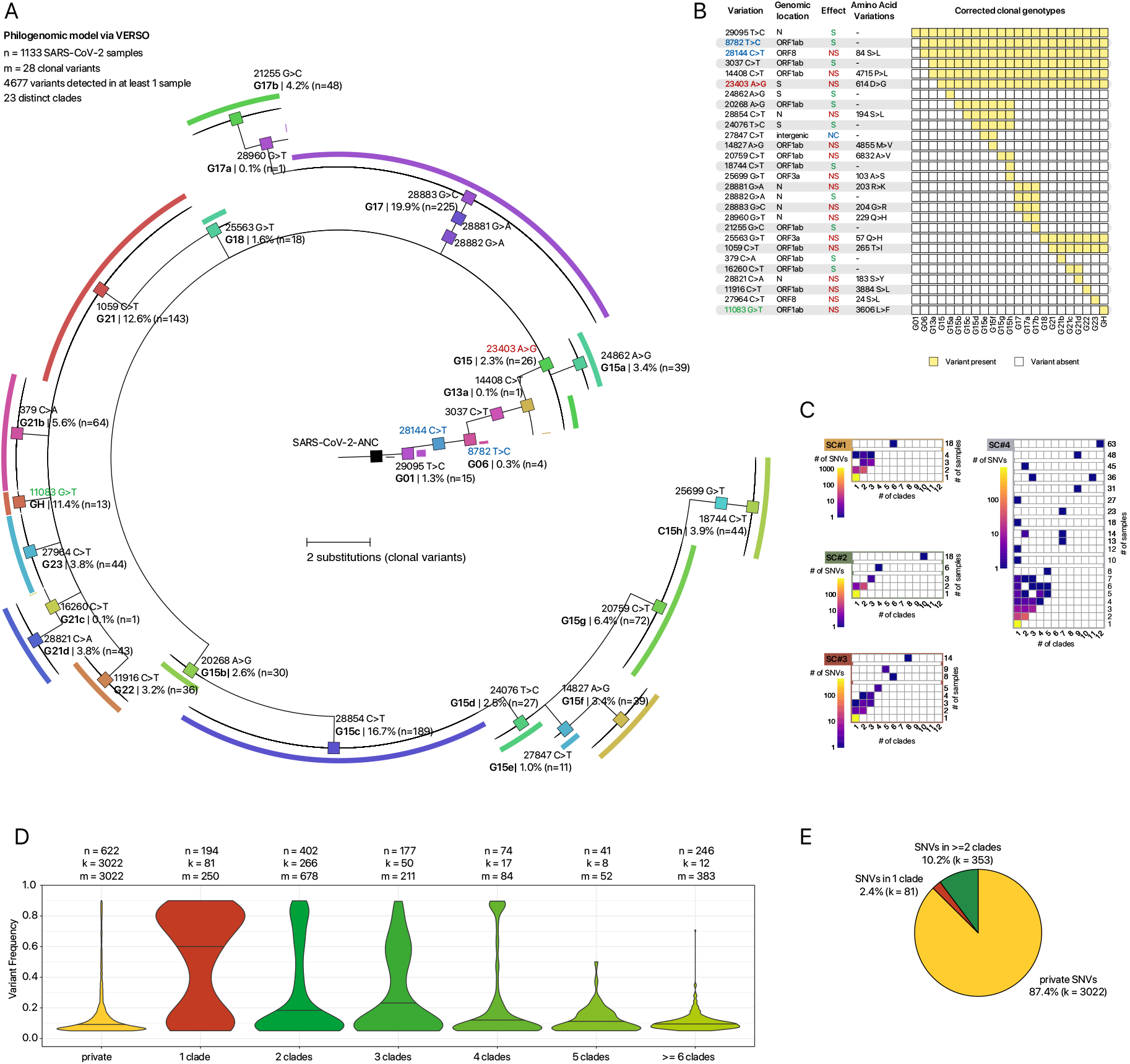
Phylogenomic model of 1133 SARS-CoV-2 samples of via VERSO – Dataset #1 (PRJNA645906) (**A**)The phylogenetic tree returned by VERSO (Ramazzotti et al., 2020) considering 28 clonal variants (VF > 0.90) detected in at least 3% of the 1133 samples of the dataset is displayed. Colors mark the 23 distinct clades identified by VERSO, which are associated to corrected clonal genotypes. Genotype labels are consistent with (Ramazzotti et al., 2020), whereas in Supplementary File S3 one can find the mapping with the lineage nomenclature proposed in Rambaut et al. (2020). Samples with identical corrected clonal genotypes are grouped in polytomies (visualization via FigTree (Rambaut, 2009)). The black colored sample represents the SARS-CoV-2-ANC reference genome. (**B**) Heatmap returning the composition of the 23 corrected clonal genotypes returned by VERSO. Clonal SNVs are annotated with mapping on ORFs, synonymous (S), nonsynonymous (NS) and non-coding (NC) states, and related amino acid substitutions. Variants g.8782T>C (*ORF1ab*, synonymous) and g.28144C>T (*ORF8*, p.84S>L) are colored in blue, variant g.23403 A>G (S, p.614 D>G) in red, homoplastic variant g.11083G>T (*ORF1ab*, p.3606L>F) in green. (**C**) Heatmaps displaying the count of minor variants with respect to the number of clades and samples in which they are found, grouped by signature-based cluster (e.g., at row 3 and column 5, the color represents the number of SNVs found in 3 clades and 5 samples). (**D**) Violin plots returning the VF distribution of all minor variants, with respect to the number of clades in which they are found (the first violin plot is associated to variants privately detected in single samples). n returns the number of samples, k the number of distinct SNVs, m the number of non-zero entries of the VF matrix. (**E**) Pie-chart returning the proportion of minor variants privately detected in single samples, detected in multiple samples of the same clade and in multiple samples of independent clades.

Interestingly, SNV g.29095T>C (mapped on ORF N, synonymous) appears to be the earliest evolutionary event from reference genome SARS-CoV-2-ANC (Ramazzotti et al., 2020). All downstream clades belong to type B type (Forster et al., 2020; Tang et al., 2020), as determined by presence of mutations g.8782T>C (*ORF1ab*, synonymous) and g.28144C>T (*ORF1ab*, p.84S>L). Importantly, we note that variant g.23403A>G (S, p.614D>G), whose correlation with viral transmissibility was investigated in depth (Lokman et al., 2020; Daniloski et al., 2020; SKorber et al., 2020; Zhou et al., 2020a; Grubaugh et al., 2020; Plante et al., 2020), is found in 18 clades, which include 1113 samples of the dataset.

In addition, the model unveils the presence of a number of *homoplasies*, as a few clonal variants are observed in independent clades and, especially, mutation g.11083G>T (*ORF1ab*, p.3606L>F), which was investigated in a number of recent studies on SARS-CoV-2 evolution (van Dorp et al., 2020; Ramazzotti et al., 2020), and is observed in 42 samples and 8 distinct clades. One might hypothesize that such SNVs have spontaneously emerged in unrelated samples and were selected either due to some functional advantage or, alternatively to the combination of founder and stochastic effects involved in variant transmission during infections, which might lead certain minor SNVs transiting to clonality in the population (see above).

As extensively discussed in Ramazzotti et al. (2020), while all the clonal variants of a host are most likely transmitted during an infection, the extent of transmission of minor variants is still baffling and is highly influenced by bottlenecks, founder effects and stochasticity (Gutierrez et al., 2012; Domingo et al., 2012). Simultaneous infections of the same host from multiple individuals harboring distinct viral lineages (also named *superinfections*), might in principle affect variant clonality, yet their occurrence is extremely rare (Lythgoe et al., 2020). For this reason, we quantified the number of minor variants (VF ≤ 90%):

1. privately detected in single samples, and which are most likely spontaneously emerged via host-related mutational processes;
2. found in multiple samples of the same clade, which might be either (*a*) spontaneously emerged or (*b*) transferred from other hosts via infection chains;
3. observed in multiple samples of independent clades (i.e., *homoplasies*), and which might be due to (*a*) positive selection of the variants due to some functional advantage, in a scenario of parallel/convergent evolution, (*b*) mutational hotspots, i.e., SVNs falling in mutation-prone sites or regions of the viral genome, (*c*) phantom mutations due to sequencing artifacts (Bandelt et al., 2002), (*d*) complex transmission dynamics involving founder effects and stochasticity, which may allow certain minor variants to transit to clonality, eventually leading to a clonal genotype transmutation (see above).

In our case, we observe that 87.4% of minor variants are observed as private of single samples, 2.4% in multiple samples of the same clade and 10.2% are detected in samples belonging to distinct clades (Fig. 5E). Importantly, significantly different VF distributions are observed and, especially, an approximately monotonic decrease of the median VF is detected with respect to the number of clades in which minor variants are found (Fig. 5D). Important conclusions can be drawn from these results.

Apparently, the large majority of minor SVNs spontaneously emerges in single samples, likely due to signature-based mutational processes. Yet, the VF distribution of private minor SNVs suggests that, as expected, most of such variants are indeed purified in the population.

Accordingly, the hypothesis of transmission of minor variants during infections is supported by the significantly larger VF of (the fewer) minor variants found in multiple samples of the same clade, as this effect is most likely due to transmission bottleneck effects (Gutierrez et al., 2012; Domingo et al., 2012).

In addition, the progressively smaller VF of minor variants observed in samples of independent clades, and which are likely more distant in the infection chain, hints at the noteworthy presence of mutational hotspots and of phantom mutations related to sequencing artifacts (Bandelt et al., 2002). In all scenarios, the presence of positively selected variant cannot be excluded, but requires ad hoc investigations.

Interestingly, one can refine the analysis by focusing on distinct signature-based clusters, for instance, by pinpointing variants likely related to mutational hotspots or phantom mutations: see, e.g., variant g.8651A>C (*ORF1ab*, p.2796M>L) which is observed in 63 samples and 12 clades (Figure 5C).

### Validation – Datasets #2 – 5

We employed 4 independent datasets (NCBI BioProjects: `PRJNA625551, PRJNA633948, PRJNA636748` and `PRJNA647529`; see Methods for details), to validate the presence of the discovered mutational signatures. Specifically, we performed signature assignment with respect to the discovered signatures on 141, 23,17 and 14 high-quality samples showing ? 6 always minor variants in each dataset, respectively.

Three signature-based clusters are found for all datasets and explain more that 97% of the variance in all cases, with highly significat p-values (see Fig 6). Such clusters are related to combinations of signatures consistently to the analysis presented in the text and display alike distributions of minor SVNs (see Supplementary Fig. S6).

**Figure 6:**
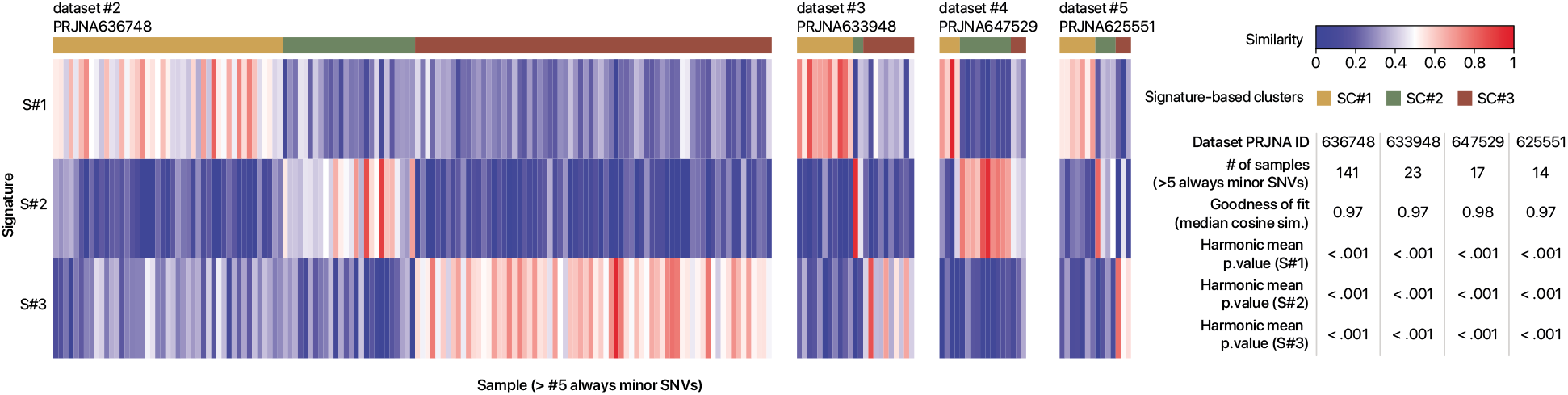
Validation on Datasets #2 – 5 (PRJNA: 636748, 633948, 647529, 625551). Heatmap returning the clustering of 141, 23, 17 and 14 samples of Datasets #2 — 5 with ≥ 6 always minor variants (≈ 13% of the dataset), computed via k-means on the low-rank latent NMF matrix on the three signatures discovered on Dataset #1 (see Methods). The goodness of fit in terms of median cosine similarity between observations and predictions, and the harmonic mean p-value of the one-sided Mann-Whitney U test on bootstrap re-sampling, are shown for all signatures, see Methods).

These important results prove the generality of our findings and strongly supports the hypothesis of distinct mutational processes active in distinct groups of samples.

## Discussion

Standard (phylo)genomic analyses of viral consensus sequences might miss useful information to investigate the elusive mechanisms of viral evolution *within hosts* and of transmission *among hosts*. In this respect, raw sequencing data of viral samples can be effectively employed to deliver a high-resolution picture of *intra-host heterogeneity*, which might underlie different clinical outcomes and affect the efficacy of anti-viral therapies. This aspect is vital especially during the critical phases of an outbreak, as experimental hypotheses are urgently needed to deliver effective prognostic, diagnostic and therapeutic strategies for infected patients.

We here presented one of the largest up-to-date quantitative analyses of intra-host genomic diversity of SARS-CoV-2, which revealed that the large majority of samples present a complex genomic composition, likely due to the interplay between host-related mutational processes and transmission dynamics.

In particular, we here proved the existence of mutually exclusive *viral mutational signatures*, i.e., nucleotide substitution patterns, which show that different hosts respond to SARS-CoV-2 infections in different ways, likely ruled by APOBEC, ROS or ADAR-related processes.

The corrected-for-signatures *dS/dN* analysis shows that such numerous low-frequency variants tend to be purified in the population whereas, conversely, a certain number of variants appear to consolidate. In particular, due to the still obscure combination of bottleneck effects and selection phenomena, certain variants appear to transit to clonality in the population, eventually leading to the definition of new clonal genotypes. Once become clonal, mutations tend to accumulate in the population, as proven by a statistically significant increase of genomic diversity, and might be used to reconstruct robust models of viral evolution via standard phylogenetic approaches.

The analysis of homoplasies, i.e., minor variants shared across distinct clades and unlikely due to infection events, demonstrate that a high number of mutations can independently emerge in multiple samples, due to mutational hotspots often related to signatures or, possibly, to positive (functional) selection. In addition, the relatively higher VF of minor variants shared by multiple samples of the same clades supports the hypothesis of transmission during infections.

To conclude, we advocate the release of a larger number of raw sequencing datasets, especially in combination with clinical data, in order to investigate the relation among the discovered host-specific processes and clinical outcome.

### Limitations of the Study

#### Reference genome

Different reference genomes have been employed for variant calling in the investigation of the origin and evolution of SARS-CoV-2. For instance, sequence `EPI_ISL_405839` was used, e.g., in Bastola et al. (2020) and sequence `EPI_ISL_402125`, e.g., in Andersen et al. (2020). As detailed in the Methods section, here we employed as reference the sequence SARS-CoV-2-ANC, which was identified in Ramazzotti et al. (2020) as a likely ancestral SARS-CoV-2 genome. Clearly, the use of different, albeit mostly overlapping, reference genomes can influence downstream analyses and, especially, the inference of the first evolutionary steps of the phylogenomic model, which should be therefore considered with caution. However, in our specific case, the employment of any of such reference genomes does not impact the identification and characterization of mutational signatures, since the SNVs that distinguish such sequences are found as clonal in at least one sample of the dataset and, accordingly, are excluded from the analysis.

#### Quasispecies composition

As discussed in the Introduction section, the analysis of raw sequencing data might be used to characterize the quasispecies architecture of single samples. To this end, a plethora of sophisticated computational methods for the characterization of the quasispecies composition of single samples is available, e.g., (Prosperi and Salemi, 2012; Giallonardo et al., 2014; Töpfer et al., 2014; Barik et al., 2018) and was recently reviewed in Knyazev et al. (2020). In the phylogenomic analysis included in this work, we decided to restrict the analysis on clonal variants that, by definition, are present in most of (or all) the quasispecies of a given sample. This allows us to provide a coarse-grained picture of the main steps of SARS-CoV-2 evolution and, at the same time, to investigate the possible transmission of minor variants, which are related to scarcely prevalent and rare quasispecies. It would be worth investigating how the combination of more sophisticated methods for quasispecies deconvolution and of our approach for mutational signatures analysis may improve the overall comprehension of SARS-CoV-2 diversity, adaptability and evolution.

#### Dataset quality

It was recently noted that some currently available SARS-CoV-2 datasets might present quality issues, especially with respect to low-frequency variants (De Maio et al., 2020). For this reason, the results of any computational pipeline should be, in principle, validated on datasets for which the ground truth is known. In our case, and given the current shortage of SARS-CoV-2 benchmark datasets, we decided to validate the discovery and characterization of the mutational signatures on 4 different datasets, generated from independent laboratories worldwide, so to ensure the generality of the results obtained via our framework (see the Validation Section).

#### Indels

The evolution of SARS-CoV-2 is characterized by the presence of a significant number of insertions and deletions (Koyama et al., 2020), which are being catalogued via COV-Glue (Singer et al., 2020). In this work, we focused on the analysis of single-nucleotide variants, as this allows us to discover and characterize statistically significant host-related mutational signatures. Despite being beyond of the scope of the current work, it might be worth investigating the origination, evolution and transmission of indels as well.

## Methods

### Datasets

#### Dataset #1

We analyzed a dataset comprising 1188 samples from NCBI BioProject with accession number PRJNA645906. For all samples, Illumina AMPLICON sequencing high-coverage raw data are provided; all patients were located in California, United States. Within this dataset, we considered for our analyses, 1133 high-quality samples having coverage ≥ 20 in at least 75% of the virus genome.

#### Datasets #2 — 5 (validation)

We considered 4 additional datasets for validation, NCBI BioProject with accession numbers `PRJNA625551` (United States, 272 AMPLICON samples), `PRJNA633948` (Australia, 203 AMPLICON samples), `PRJNA636748` (South Africa, 408 AMPLICON samples), `PRJNA647529` (Israel, 212 AMPLICON samples). We applied the same QC filters to these datasets (coverage ≥ 20 in at least 75% of virus genome), to obtain four validation set including a total of 953 samples.

#### SNVs calling

For all datasets, we downloaded SRA files and converted them to FASTQ files using SRA toolkit. Following (Ramazzotti et al., 2020), we used Trimmomatic (version 0.39) to remove positions at low quality from the RNA sequences, using the following settings: LEADING:20 TRAILING:20 SLIDINGWINDOW:4:20 MINLEN:40.

We used bwa mem (version 0.7.17) to map reads to the reference genome SARS-CoV-2-ANC, which was recently released in Ramazzotti et al. (2020). SARS-CoV-2-ANC is identical to `EPI_ISL_405839` (Bastola et al., 2020) and `EPI_ISL_402125` (Andersen et al., 2020) reference genomes on 29865 (out of 29870) genome locations (99.9%), includes the polyA tail of the latter genome (33 bases), and has haplotype TCTCT at locations 8782, 9561, 15607, 28144 and 29095, as observed in both the Bat-CoV-RaTG13 (sequence `EPI_ISL_402131`) (Zhou et al., 2020b) and Pangolin-CoV (sequence `EPI_ISL_410721`) (Andersen et al., 2020; Xiao et al., 2020) genomes.

We then generated sorted BAM files from bwa mem results with SAMtools (version 1.6) and removed duplicates with Picard (version 2.22.1). Variant calling was performed generating mpileup files with SAMtools and then using VarScan (min-var-freq parameter set to 0.01) (Koboldt et al., 2012).

We finally verified the absence of any possible bias in the detection of minor variants due to sequencing artifacts. As one can see in Supplementary Fig. S1, no correlation between the number of SNVs and both total coverage and the median coverage is observed (R^2^ < 0.04 in both cases), which proves the good quality of the calls.

### Signatures analysis

The analysis was performed on always minor variants (VF > 5% and ≤ 90% in all samples in which they are detected), in order to ensure that the considered variants are not due to transmission, but are likely emerged in the host. In such way, we could associate to each discovered signature a mechanism causing variants in the viral genome related to the specific host.

Signatures decomposition was formulated as a Non-negative Matrix Factorization problem (NMF) (Brunet et al., 2004). Given *n* samples, *r* possible substitution classes (e.g., C>T:G>A) and s signatures, we can define the following objects:

- the input data matrix **D**, a *n* × *r* dimensional matrix, where every element *d_i,j_* represents the number of SNVs with substitution class *j* in the *i^th^* sample. Note that 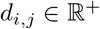;
- the low-rank latent NMF matrix **A**, a *n* × *s* dimensional matrix, where every element *a_i,j_* represents the linear combination coefficient of signature *j* in sample *i* (also *exposure* of the *i*^th^ sample to signature *j* (Alexandrov et al., 2013)). Note that 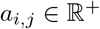;
- the signature (or basis) matrix **B**, a *s* × *r* dimensional matrix, where every row is a categorical distribution of all substitution classes in each signature. For this matrix we assume that every row must sum up to 1, then *b_i,j_* ∈ {0,1} and 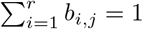.

In particular, we here considered 6 substitution classes (*r* = 6) by merging equivalent substitution types, namely G>T:C>A, G>C:C>G, G>A:C>T, A>T:T>A, A>G:T>C and A>C:T>G.

#### Problem 1

*(Signature decomposition): Given the data matrix* **D**, *we aim at finding the NMF latent matrix* **A** *and the signature matrix* **B**, *such that* ║**D** – **A** · **B**║_2_ *is minimum*.

To solve the stated problem, we here performed a total of 100 independent NMF runs with standard update (Brunet et al., 2004), for solutions at ranks varying from 1 to 6, where initial solutions were randomly initialized; for each run a total of 20 iterations were performed where signatures and their assignments to samples were iteratively estimated by non-negative least squares (Chen and Plemmons, 2010). The final solution was constructed as the consensus of the 100 runs (Brunet et al., 2004).

We then employed multiple state-of-the-art approaches to assess the optimal rank (optimal number of signatures s) for the NMF decomposition. We first assess the stability of NMF results over the 100 runs, with the idea that stable solutions are preferable to unstable ones; to this extent, we computed the average Cophenetic correlation coefficient (Brunet et al., 2004), which showed high stability (0.998) at rank equal to 3 (see Supplementary Fig. S3A). Furthermore, we also evaluated the goodness of fit of NMF solutions at different ranks, with rank equals to 3 being able to explain 96.54% of variance in the data (see Supplementary Fig. S3B); finally, we report as Supplementary Fig. s3C the average cosine similarity between observations and predictions by NMF with rank equals to 3, showing a plateau with correlation equals 0.973 (Lal et al., 2020). All of this supports 3 as the optimal rank for our decomposition problem and the presence of 3 distinct mutational signatures in our data.

#### Identification of signature-based clusters

In order to identify clusters of samples possibly affected in different proportions by the discovered mutational signatures, we considered the low-rank latent NMF matrix **A** defined above.

Specifically, we first normalized **A** such that each row of the matrix sums up to 1 and then computed the euclidean distance among each pair of samples. We next performed Principal Component Analysis (PCA) on the distance matrix to estimate the optimal number of clusters present in our data. In detail, the analysis of the eigenvalues of the distance matrix shows that 3 components explain > 99% of the variance, followed by a plateau. Accordingly, we performed k-means clustering with *k* = 3 on the normalized **A** matrix to discover the signature-based clusters.

#### Assessment of signatures significance

We assessed the statistical significance of samples exposure to signatures by bootstrap. Namely, for each sample we performed 1000 bootstrap re-sampling from their observed variants cumulative distributions and assigned signatures to each bootstrap dataset in order to obtain 1000 independent assignments for each sample. Then, for each signature we computed a p-value by Mann-Whitney U test to verify the hypothesis that such signature was contributing to more than 5% variants (one-sided test). A p-value assessing the significance of each signature was computed as the harmonic mean of the Mann-Whitney U test p-values (Wilson, 2019).

### Corrected-for-signatures dN/dS analysis

In order to quantify the selection pressure in coding regions of SARS-CoV-2, we employed *dN/dS* analysis, which assesses and compares non-synonymous to synonymous substitution rates. In its standard version, this analysis assumes uniform nucleotide substitution probabilities across the genome; however, this hypothesis might not hold if different mutational processes are active with biases over a subset of substitutions (e.g., non-uniform distribution might be observed across signature-based clusters). If this bias is not taken in account, it may lead to erroneous estimation of the *dN/dS* ratio (Van den Eynden and Larsson, 2017).

For this reason, since we discovered the existence of different host-related mutational processes (i.e., the mutational signatures) that are strongly biased toward specific substitutions, we developed a corrected-for-signatures *dN/dS* ratio analysis, as proposed in a different context in Van den Eynden and Larsson (2017). Specifically, given the *i^th^* sliding window of the coding region comprising *l* bases and considering the *f^th^* signature-based cluster, the corrected *dN/dS* ratio (for signature-cluster *f* and sliding window *i*) is given by:

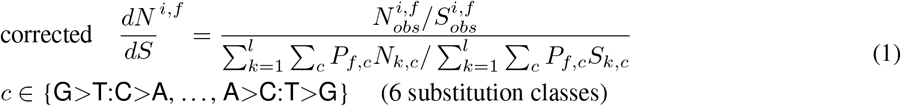

where, 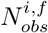 and 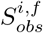, are the numbers of non-synonymous and synonymous substitutions detected in the window *i* in at least one sample of signature-based cluster *f, P_f,c_* is the probability of substitution class *c* in signature-based cluster *f* (computed with respect to the categorical normalized cumulative VF distribution of all substitution classes in that cluster), and *N_k,c_* (*S_k,c_*) is equal to 1 if class c identifies an admissible non-synonymous (synonymous) substitution in the *k^th^* position of the window *i*, 0 otherwise.

### Phylogenetic model from clonal variant profiles via VERSO

VERSO is a 2-step computational framework for the characterization of viral evolution from raw sequencing data introduced in Ramazzotti et al. (2020). In particular, VERSO STEP #1 is a probabilistic noise-tolerant approach that processes binarized clonal variant profiles to deliver a robust phylogenetic model also in condition of sampling limitations and sequencing issues (for further details please refer to the related article).

In this work, we followed the guidelines proposed in Ramazzotti et al. (2020) and applied VERSO STEP #1 to the binarized mutational profiles of clonal mutations to reconstruct a phylogenetic model of the SARS-CoV-2 samples included in our dataset, and to investigate the possible presence of homoplasies of minor variants. In our analysis, we considered only clonal variants (VF > 90%) detected in at least 3% of the samples of the dataset (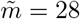 clonal variants on *n* = 1133 samples). The method was executed with 1 million MCMC iterations and returns 23 distinct clades associated to corrected clonal genotypes returned by the method. The stratification of samples in clades was employed in the analysis of homoplasies of minor variants.

We note that several widely used alternative approaches for phylogeny reconstruction from consensus sequences are available, among which, e.g., IQ-TREE (Nguyen et al., 2015), the algorithm included in the Nextstrain-Augur pipeline (Hadfield et al., 2018), BEAST 2 (Bouckaert et al., 2019) and MrBayes (Ronquist et al., 2012). In Supplementary Fig. S5 the phylogenetic model returned via MrBayes (Ronquist et al., 2012) on Dataset #1 is displayed (the method was executed with 10 million MCMC iterations and default parameters), showing consistent results with our analysis, as proven by the Adjusted Rand Index (ARI) (Santos and Embrechts, 2009) between sample partitionings (ARI = 0.76).

## Software availability

The source code used to replicate all the analyses is available at this link: https://github.com/BIMIB-DISCo/SARS-CoV-2-IHMV.

VERSO can be downloaded at this link: https://github.com/BIMIB-DISCo/VERSO.

## Authors contributions

A.G., D.M., F.A., R.P. and D.R. designed and developed the study. A.G, D.M., F.A. and D.R defined, implemented and executed the computational analyses. A.G., D.M., F.A., R.P. and D.R. analyzed the data and interpreted the results. A.G., D.R. and R.P. supervised the study. All authors wrote the manuscript, discussed the results, and commented on the manuscript.

## Acknowledgments

This work was partially supported by the Elixir Italian Chapter and the SysBioNet project, a Ministero dell’Istruzione, dell’Università e della Ricerca initiative for the Italian Roadmap of European Strategy Forum on Research Infrastructures and by the AIRC-IG grant 22082. We thank Marco Antoniotti, Giulio Caravagna and Chiara Damiani for helpful discussions.

