## Supplementary Information for "Mutational signatures and heterogeneous host response revealed via large-scale characterization of SARS-CoV-2 genomic diversity"

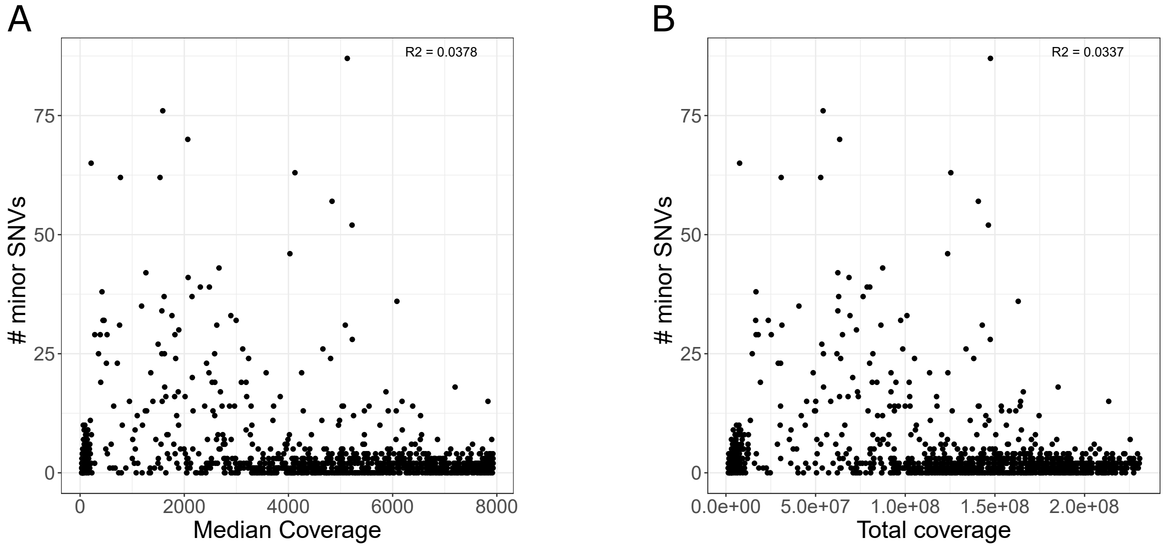

Figure S1: **Quality check – Dataset** #1 `PRJNA645906`. (A) Scatter-plot returning for each sample of the cohort the number of minor SNVs (variant frequency VF $\leq 90\%$) and the median coverage. $R^2$ coefficient is shown in the upper-right corner. (B) Scatter-plot returning for each sample of the cohort the number of minor SNVs and the total coverage.

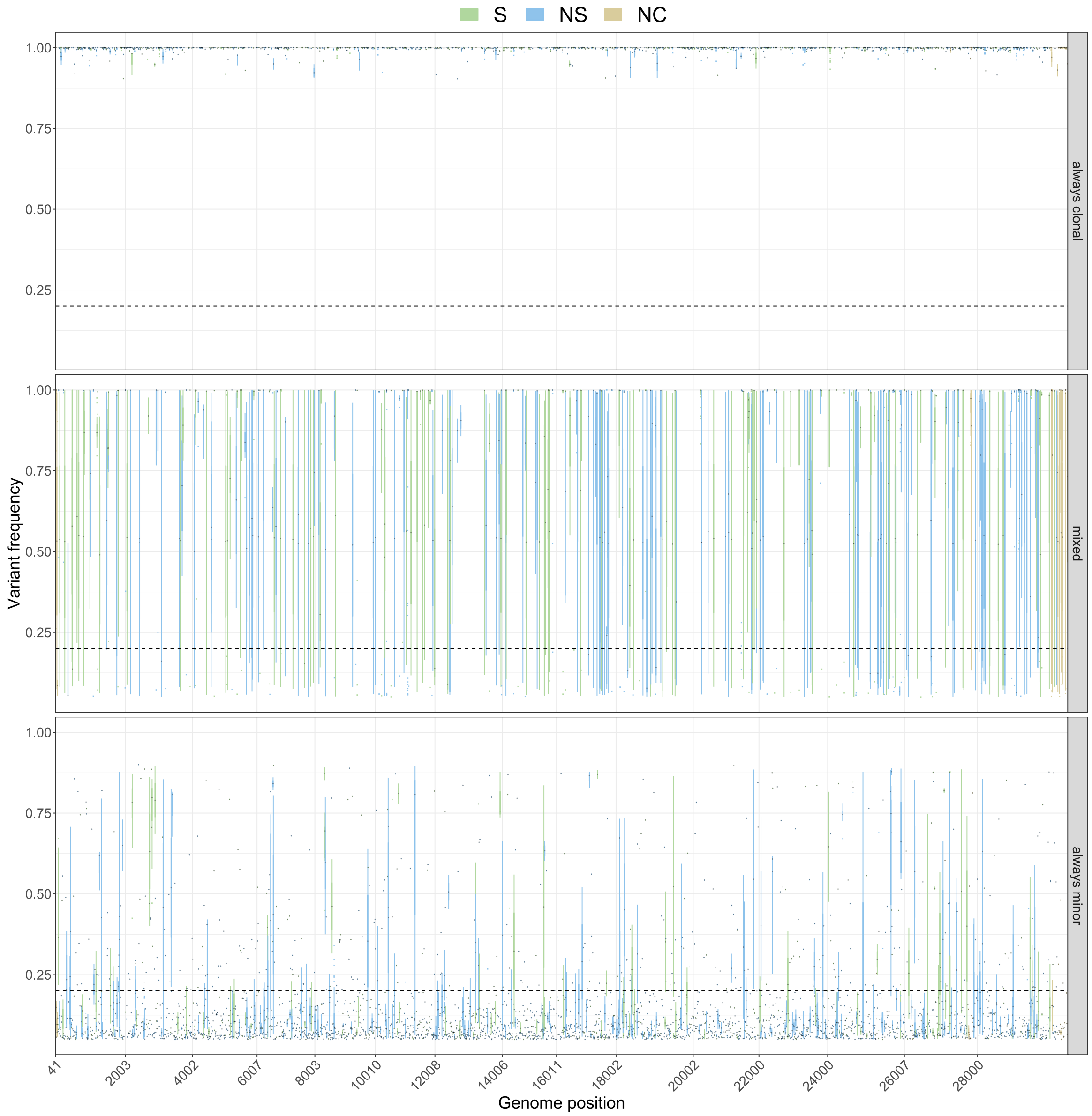

Figure S2: **Distribution of SNVs detected on SARS-CoV-2 genome – Dataset #1 `PRJNA645906`.** Box-plots returning the VF distribution of all SNVs detected in all samples, colored according synonymous, non-synonymous and non-coding state and grouped according to SNV category.

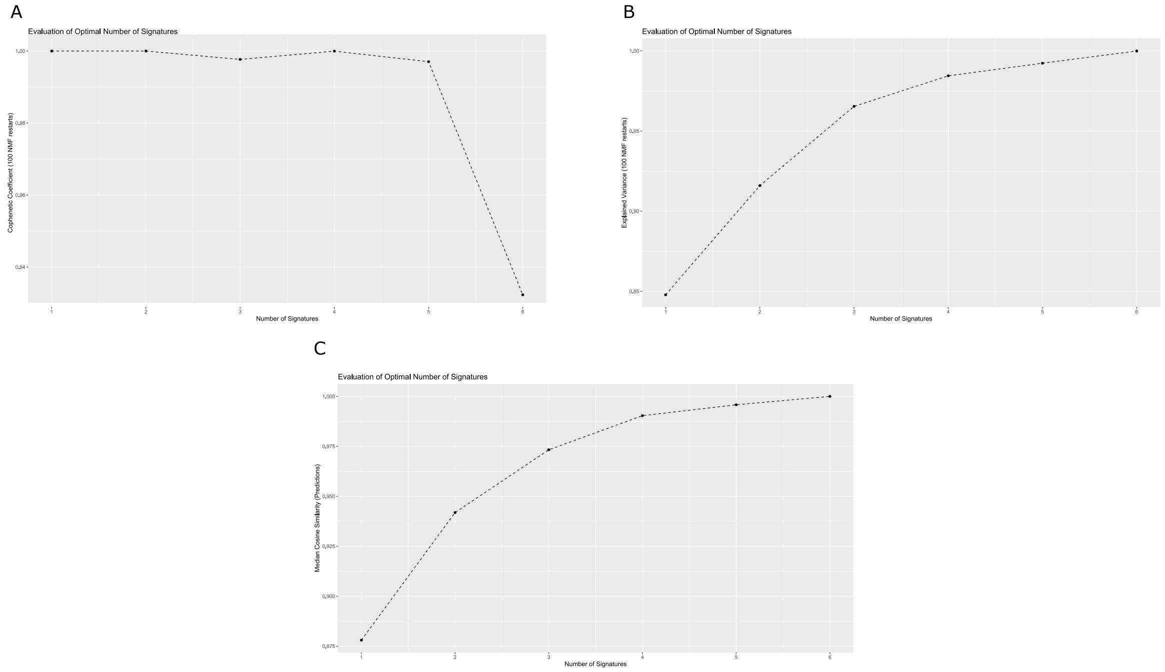

Figure S3: **Signature analysis: Cophenetic correlation coefficient, dispersion coefficient, explained variance, goodness of fit – Dataset** #1 `PRJNA645906`. (A) Explained variance, (B) average Cophenetic correlation coefficient with respect to Non-negative Matrix Factorization (NMF) rank, in the range $1-6$. 1000 NMF restarts comprising 20 iterations were performed. A sharp drop is observed between 3 and 4, suggesting that the optimal rank (i.e., the number of signatures) is 3, (C) goodness of fit, measured as average Cosine correlation coefficient among observed profiles and predicted ones, with respect to NMF rank in the range $1-6$ (Lal et al., 2020). A plateau is observed at rank 3.

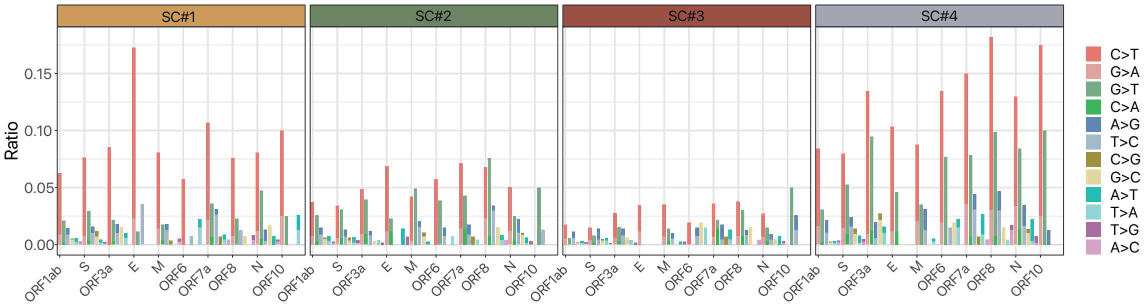

Figure S4: **Distribution of substitution types on SARS-CoV-2 ORFs – Dataset #1** `PRJNA645906`. Barplots displaying the categorical distribution of all SNVs detected in all signature-based clusters, with respect SARS-CoV-2 ORFs (normalized by reference allele count in each ORF).

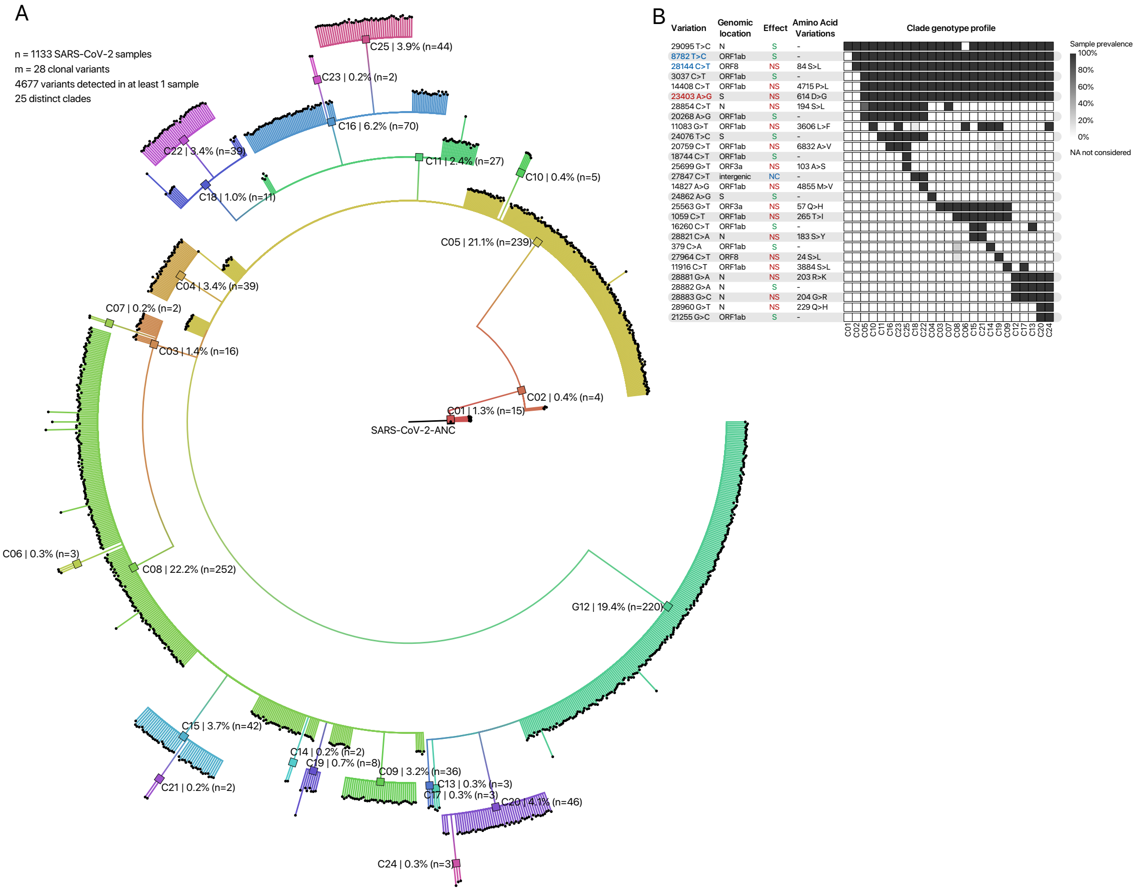

Figure S5: **Phylogenomic model of** 1133 **SARS-CoV-2 samples returned by MrBayes – Dataset** #1 PRJNA645906). **(A)** The phylogenetic tree returned by MrBayes (Ronquist et al., 2012) considering 28 clonal variants (VF > 0.90) detected in at least 3% of the 1133 samples of the dataset is displayed. Colors mark the 25 clades identified by MrBayes (visualization via FigTree (Rambaut, 2009)). The black colored sample represents the SARS-CoV-2-ANC reference genome. **(B)** Heatmap returning the fraction of samples of any clade in which a specific clonal variant is observed. Clonal SNVs are annotated with mapping on ORFs, synonymous (S), nonsynonymous (NS) and non-coding (NC) states, and related amino acid substitutions. Variants g.8782T>C (*ORF1ab*, synonymous) and g.28144C>T (*ORF8*, p.84S>L) are colored in blue, whereas variant g.23403 A>G (*S*, p.614 D>G) is colored in red. The results are consistent with those obtained by VERSO STEP #1 (Figure 5 of the main text), as proven by the Adjusted Rand Index (ARI) (Santos and Embrechts, 2009) between sample partitionings (ARI = 0.76)

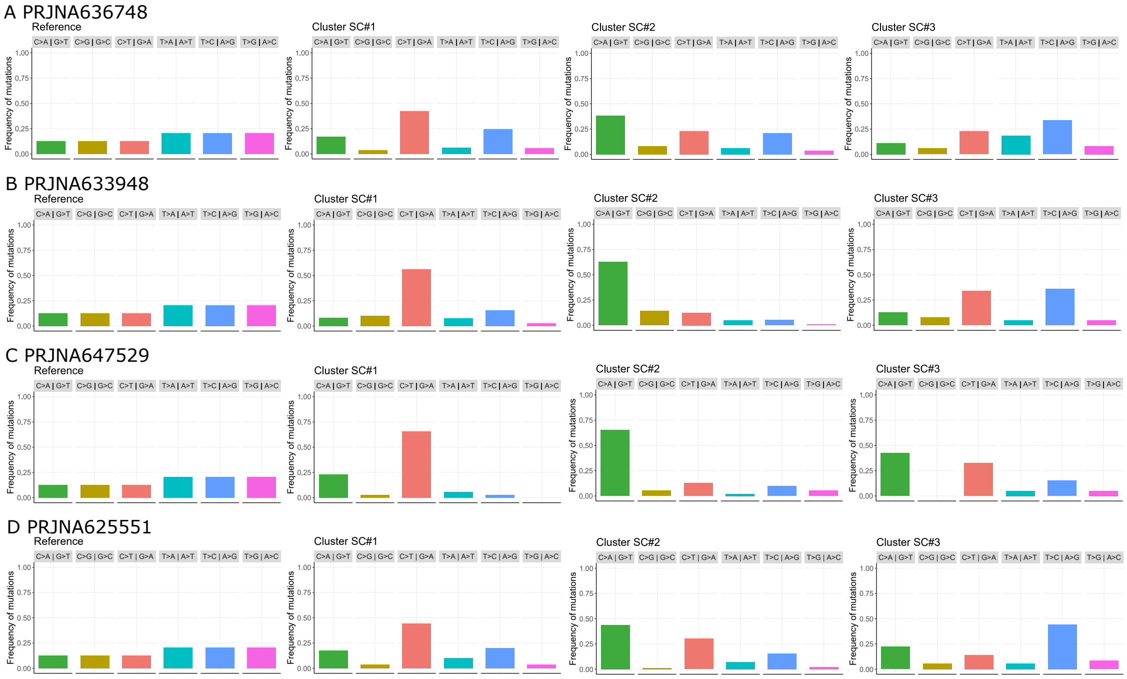

Figure S6: **Categorical substitution type distribution – Validation datasets** $\#2, 3, 4, 5$ (`PRJNA: 636748, 633948, 647529, 625551`). Categorical normalized cumulative distribution of all SNVs detected in each signature-based cluster of the validation datasets, with respect to 6 substitution classes.
